# Chemoresistance to additive PARP/PI3K dual inhibition in triple-negative breast cancer cell lines is associated with adaptive stem cell-like prevalence

**DOI:** 10.1101/2024.04.28.591568

**Authors:** Paula E. Petrella, Jason W. Chen, Gabrielle O. Ravelo, Benjamin D. Cosgrove

## Abstract

Cancer stem-like cells (CSCs) are posited to exhibit specialized oncogenic capacity to drive malignancies. CSCs are distinguished by enhanced hallmarks of cancer, including apoptosis avoidance, phenotypic plasticity and aberrant growth pathway signaling. Standard-of-care chemotherapies targeted to rapidly cycling cells routinely fail to eliminate this resistant subpopulation, leading to disease recurrence and metastasis. Triple-negative breast cancer (TNBC), a highly aggressive subtype of breast cancer, is enriched for tumor-propagating CD44^+^/CD24^−/low^ CSCs, which are poorly ablated by chemotherapeutics and are associated with poor prognosis. CD44 governs sustained PI3K signaling in breast cancer, which is essential for CSC maintenance. PI3K inhibition can elicit DNA damage and down-regulate BRCA1 expression, which in turn enhance the synthetic lethality of PARP inhibitors. Here, we examined a dual chemotherapeutic approach targeting these pathways by combining a pan-PI3K inhibitor (Buparlisib) and a PARP1 inhibitor (Olaparib) on a panel of TNBC cell lines with distinct mutational profiles and proportions of CSCs. We observed differential sensitivity to this dual inhibition strategy and varying cellular stress and resistance responses across eight TNBC lines. The dual chemotherapeutic effect is associated with a reduction in S-phase cells, an increased in apoptotic cells and elevated expression of cleaved PARP, indicating a provoked replicative stress response. We observed that PARP/PI3K inhibition efficacy was potentiated by repeated administration in some TNBC lines and identified critical treatment schedules, which further potentiated the dual chemotherapeutic effect. Dual inhibition induced small but significant increases in CSC relative abundance as marked by CD44^+^/CD24^−/low^ or ALDH1^+^ cells and increased stress and survival signaling in multiple TNBC cell lines, suggesting this sub-population contributes to TNBC chemoresistance. These results suggest the additive effects of PARP and PI3K inhibition against CSC phenotypes may be enhanced by temporally-staged administration in TNBC cells.

## Introduction

Breast cancer (BC) is the most prevalent type of cancer diagnosed in women, and the second most lethal (Siegel, Miller, Wagle, & Jemal, 2023). The triple-negative BC subtype (TNBC), traditionally defined as estrogen and progesterone receptor negative and lacking in HER2 receptor over-expression (ER^−^/PR^−^/HER2^−^, respectively), is observed in 10% of new BC cases (“Surveillance Research Program, National Cancer Institute,” 2023), and is challenging to treat with the highest metastatic capacity and shortest time to relapse (Cheng et al., 2016; Dent et al., 2007; Echeverria et al., 2018; Oskarsson, Batlle, & Massague, 2014). TNBC’s dearth of therapeutic targets and enrichment with chemo-resistant cancer stem cells (CSCs) (Fultang, Chakraborty, & Peethambaran, 2021; He et al., 2019; H. Li et al., 2013; O’Conor, Chen, Gonzalez, Cao, & Peng, 2018) limit current treatment options. Since the discovery of tumor-initiating stem cells in leukemia in 1994 (Lapidot et al., 1994), the CSC hypothesis has evolved to contemplate a stem-like subpopulation of chemo-resistant cells, which exhibit specialized capacity of initiating and sustaining tumors. Phenotypic survival advantages employed by CSCs to maintain chemo-resistance include robust DNA repair, alteration of therapeutic targets, dormancy, apoptosis avoidance and increased expression of drug efflux pumps (Batlle & Clevers, 2017; Dean, Fojo, & Bates, 2005; Hanahan & Weinberg, 2011; Kreso & Dick, 2014; Oren et al., 2021; Pattabiraman & Weinberg, 2014; Visvader & Lindeman, 2008; Wicha, Liu, & Dontu, 2006).

Human breast cancer stem-like cells (BCSCs) have been characterized as a CD44^+^/CD24^−/low^ sub-population. Notably, Al-Hajj, et al. (Al-Hajj, Wicha, Benito-Hernandez, Morrison, & Clarke, 2003) showed that as few as 100 CD44^+^/CD24^−/low^ BC cells were tumorigenic in NOD-SCID mice throughout serial implantation (Chekhun et al., 2017; Idowu et al., 2012; Li et al., 2017). The CD44 cell surface receptor is involved in cell signaling, attachment and migration and, upon binding its primary ligand hyaluronic acid, an integral component of the extracellular matrix (Chen, Zhao, Karnad, & Freeman, 2018; Senbanjo & Chellaiah, 2017), has been shown to upregulate drug resistance and anti-apoptotic proteins in BC (Bourguignon, Spevak, Wong, Xia, & Gilad, 2009). The CD24 receptor is regarded as a mark of differentiation (Jaggupilli & Elkord, 2012; Shipitsin et al., 2007) in the mammary epithelium and its suppressed expression may indicate a stem-like BC cell capable of phenotypic plasticity. Expression of the detoxifying enzyme ALDH1 has also been associated with BCSCs (Ginestier et al., 2007) and may play a role in BC chemo-resistance (Al-Shamma, Zaher, Hersi, Abu Jayab, & Omar, 2023). Critically, both ALDH1 expression and a CD44^+^/CD24^−/low^ phenotype have been correlated with metastatic potential and a poor prognosis in BC patients (Horimoto et al., 2016; Idowu et al., 2012; Ma et al., 2017; Sheridan et al., 2006; Zhong et al., 2014).

Traditional chemotherapies target vulnerabilities common to most cells in a tumor, often by targeting quickly cycling cells, but are less effective at eradicating rare, slow-cycling CSCs, leading to disease recurrence and progression to metastasis (Wicha et al., 2006). Failure of traditional chemotherapeutics in BC may be explained by an increase in the proportion of CD44^+^/CD24^−^ and ALDH1^+^ BCSCs in human samples after treatment with docetaxel (Creighton et al., 2009); doxorubicin and either docetaxel or cyclophosphamide (Lee et al., 2011); or vincristine (Ghanbari, Azadbakht, Vesi-Raygani, & Khazaei, 2016), and the induction of CD44^+^/CD24^low^ BCSCs by metabolic stress in TNBC (Jaggupilli et al., 2022). Indeed, extended treatment with doxorubicin enriches the CD44^+^/CD24^−^ fraction in BC (Calcagno et al., 2010). Conversely, knockdown or downregulation of CD44 in BC cells increases sensitivity to conventional therapies (Pham et al., 2011; Van Phuc et al., 2011), and CD24 over-expression enhances TNBC vulnerability to doxorubicin (Deng et al., 2017). Hence, there remains a compelling need to identify targetable vulnerabilities in the BCSC sub-population that is resistant to standard chemotherapies.

We considered here whether the efficacy of combined PI3K and PARP1 inhibition in BC (Juvekar et al., 2012) would attenuate the BCSC phenotype in TNBC cell lines via mechanisms distinct from those targeted ineffectually by traditional chemotherapies. PI3K, a key lipid kinase upstream from Akt, is involved in multiple signaling pathways governing cell growth, proliferation, protein synthesis and suppression of apoptosis (Fig. 1a), and has been implicated in the maintenance of BCSCs (J. Zhou et al., 2007). Activating mutations in PI3K are common in BCs (Isakoff et al., 2005) and can contribute to chemo-resistance (Hu et al., 2021). Notably, CD44 induces sustained PI3K/Akt signaling (S. Liu & Cheng, 2017) and contributes to PI3K inhibition resistance in BC (Yang et al., 2020). PARP1 recruits DNA damage repair proteins at both single- and double-strand breaks (Ray Chaudhuri & Nussenzweig, 2017) (Fig. 1a) and is overexpressed in TNBC cells (Gilabert et al., 2014). In cells with BRCA1 mutations, which comprise 10-15% of TNBCs (Choi et al., 2023), PARP inhibition can lead to synthetic lethality (Pommier, O’Connor, & de Bono, 2016). In turn, PI3K inhibition has been found to downregulate BRCA1 expression in TNBC, inducing sensitivity to PARP inhibition (Ibrahim et al., 2012). Importantly, PARP1 is required for somatic cell emergence from quiescence (Mostocotto et al., 2014) and progression from G0 to G1 (Carbone et al., 2008), and PI3K is known to control entry into the cell cycle (Garcia, Kumar, Marques, Cortes, & Carrera, 2006), with its oncogenic hyperactivity linked to CSC stemness and induction of EMT (Madsen, 2020).

**Figure 1.**
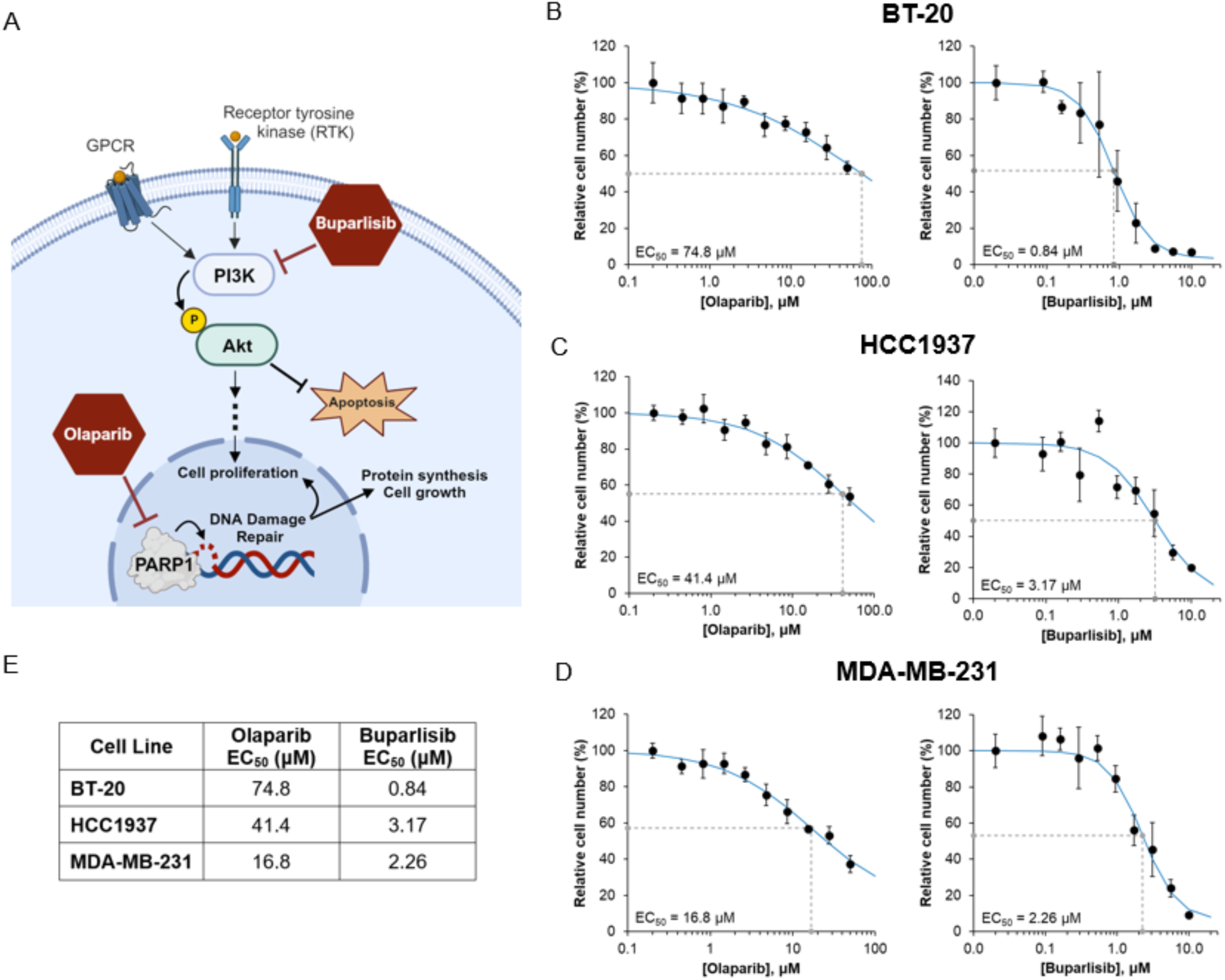
PARP and PI3K inhibition limits TNBC cell viability *in vitro*. (A) Scheme of the inhibitory effects of Olaparib and Buparlisib, small molecule PARP1 and pan-PI3K inhibitors, respectively. The lipid kinase PI3K phosphorylates Akt, activating downstream effects. PARP1, a nuclear enzyme, recruits DNA repair proteins to single- and double-stranded breaks. (B-E) Single-drug dose responses at 72 hrs for BT-20 (B), HCC1937 (C) and MDA-MB-231 (D) TNBC cell lines. Cells were treated for 72 hrs with each compound, stained with propidium iodide, and counted. Data is mean ± SD of n = 3 or more replicates and fit to a 4PL model to calculate EC_50_ values (shown in dashed lines). Single-drug EC_50_ values are reported in insets and summarized in (E).

To leverage the synergy between the pan-PI3K inhibitor (PI3Ki) Buparlisib (BUP) and Olaparib (OLA), an FDA-approved PARP1 inhibitor (PARPi) reported in BC xenograft models, we tested these small molecule drugs on TNBC cell lines harboring PI3K or BRCA1 mutations, or neither, and assessed their effects on cell survival and the proportions of CD44, CD24 and ALDH1 expressing cells. We investigated the finding by Juvekar et al. (Juvekar et al., 2016) that PI3K inhibition in BC causes cell death through nucleoside depletion, by temporally-sequenced delivery of each drug in comparison to contemporaneous treatment, and found advantages in single-drug priming. We show that combined PI3K and PARP1 inhibition increases stress-related signaling and induces apoptosis in TNBC cells with small but significant relative increases in the CD44^+^/CD24^−^ phenotype and ALDH1-expressing cells. With repeat treatment we also noted the development of cell line-specific chemoresistance. Our findings provide insight into previously uncharacterized effects of combined PARP and PI3K inhibition on TNBC cells and advances our understanding of how the inhibition of targets critical to CSC survival affect the prevalence of the stem-like subpopulation. Further study is warranted to discern optimal treatment schedules and drug concentrations based on the mutational profiles of each cell line.

## Results

### Olaparib (PARPi) and Buparlisib (PI3Ki) inhibit TNBC cell viability *in vitro*

The combination of the PARPi Olaparib and PI3Ki Buparlisib has been shown to synergize in a murine model of *BRCA1* mutation-related breast cancer (Juvekar et al., 2012), and PI3K blockade is known to sensitize *BRCA1*-proficient TNBC cell lines to PARP inhibition, implicating PI3K in DNA damage repair (Ibrahim et al., 2012) (Y. Li et al., 2021). We first tested the efficacy of these two small molecule inhibitors individually in a panel of TNBC cell lines with diverse mutational backgrounds (Table 1), including *BRCA1*-mutated, *PTEN*-deleted HCC1937; BT-20 which harbors two activating *PI3KCA* mutations; and the MDA-MB-231 metastatic cell line which is not known to carry these mutations (Fig. 1). After 72 hrs of continuous treatment over a broad range of treatment concentrations in all three lines, cells were stained with propidium iodide and counted using an automated CellProfiler pipeline (Stirling et al., 2021). Based on single-agent dose responses, the half-maximal concentration (EC_50_) was calculated using the 4PL curve fit function in Excel (Fig. 1b-e), which revealed greater efficacy of Buparlisib (PI3Ki) than Olaparib (PARPi) at low micromolar concentrations. A lower concentration of Buparlisib (EC_50_ from 0.84 to 3.17 μM) than Olaparib (EC_50_ from 16.8 to 74.8 μM) was required to achieve 50% therapeutic efficacy in all three cell lines (Fig. 1d). BT-20 cells exhibited the greatest sensitivity to PI3K inhibition, demonstrating their reliance on the two activating mutations in the kinase’s catalytic subunit (Gymnopoulos, Elsliger, & Vogt, 2007; Vasan et al., 2019), and the lowest sensitivity to PARP inhibition, as these cells express BRCA1 at a high level (Wesierska-Gadek et al., 2012). Although the HCC1937 cells harbor a BRCA1 mutation, they required a higher concentration of Olaparib than BRCA1-competent MDA-MB-231 cells to reach EC_50_, pointing to a pre-existing rewiring of the 1937 BRCA1 pathway enabling functional reliance on ATR (Yazinski et al., 2017) and RAD51 (Y. Liu et al., 2017). Their *PTEN* deletion may contribute to their relative insensitivity to PI3K inhibition, requiring the highest concentration of Buparlisib of the three cell lines to reach EC_50_. The 231 cells, lacking these mutational profiles, required the lowest concentration of Olaparib and a median concentration of Buparlisib to reach EC_50_. These distinct responses to single-inhibitor treatments expose vulnerabilities that may be exploited with combined PARP and PI3K inhibition.

**Table 1.** Model human triple-negative breast cancer cell lines. Histology, tumor source, commonly mutated genes and mutated loci in each cell line used in this study. Adapted from cellmodelpassports.sanger.ac.uk; SIB Swiss Institute of Bioinformatics’ cellosaurus.com; and ATCC.org.

| Cell Lines | Tumor Subtype | Histology | Tumor Source | Commonly Mutated Genes | Mutated Loci (cDNA / protein) |
| --- | --- | --- | --- | --- | --- |
| **BT-20** | Unclassified | Carcinoma | Primary | *CDKN2A* - homozygous<br>*PIK3CA* - heterozygous<br>*PIK3CA* – heterozygous<br>*RB1* - heterozygous<br>*TP53* - homozygous | c.1_471del471 / p.0?<br>c.1616C>G / p.Pro539Arg<br>c.3140A>G / p.His1047Arg<br>c.1544C>T / p.Pro515Leu<br>c.394A>C / p.Lys132Gln |
| **HCC1937** | Basal-like 1 | Primary ductal carcinoma | Primary | *BRCA1* - homozygous<br>*PTEN* - homozygous<br>*RB1* - homozygous<br>*TP53* - homozygous | c.5266_5267insC / p.Gln1756Profs*74<br>Deletion<br>c.2212_2325del114 / p.Thr738_Arg775del38<br>c.916C>T / p.Arg306Ter |
| **MDA-MB-231** | Mesenchymal stem-like | Adenocarcinoma | Metastasis, pleural effusion | *BRAF* - heterozygous<br>*CDKN2A* – homozygous<br>*KRAS* - heterozygous<br>*NF2* - homozygous<br>*TP53* - homozygous | c.1391G>T / p.Gly464Val<br>c.1_471del471 / p.0?<br>c.38G>A / p.Gly13Asp<br>c.691G>T / p.Glu231*<br>c.839G>A / p.Arg280Lys |

### Combined PARP and PI3K inhibition synergize at low micromolar concentrations

To examine the effects of combined PARPi and PI3Ki treatment on TNBC cell viability, we implemented a combinatorial matrix of drug concentrations to determine the most efficacious pairing (Fig. 2a-c). Cells were fixed, stained, and counted at 72 hrs as above, and data was normalized to vehicle control (VC). In testing PARPi at concentrations of 0.5 to 25 μM and PI3Ki at concentrations of 0.2 to 10 μM, we noted more potent viability restriction at multiple combinations as compared to single-agent treatments (Fig. 1). In BT-20 cells, PARPi treatment of 5.0 μM resulted in 66.9% viable cells, but in combination with 2.0 μM PI3Ki, viability was reduced to 16.1%, underscoring the acute sensitivity of BT-20 cells to PI3Ki (Fig. 2d). HCC1937 cells exhibited a similar but less drastic trend in that PARPi at 5.0 μM elicited 73.0% viability while the addition of 2.0 μM PI3Ki brought viability down to 45.8%. Treating the MDA-MB-231 cells with 5.0 μM PARPi reduced viability to 63.9% as compared to 35.9% when combined with 2.0 μM PI3Ki. (Dual-drug viability heatmaps for six additional TNBC cell lines are provided in Fig. S1.) Hence, a combinatorial approach allowed the use of a much lower concentration of PARPi than required for each cell line’s individual EC_50_, and a lower concentration of the PI3K inhibitor than two of the cell lines’ EC_50_. We elected to use the PARPi at 5.0 μM and the PI3Ki at 2.0 in subsequent experiments given the cell lines’ distinctive response signatures, and to prevent either drug from dominating the response outcomes. Taken together, these results indicate combined PARP/PI3K inhibition decreases TNBC cell viability across mutational profiles through additive dual-agent effects.

**Figure 2:**
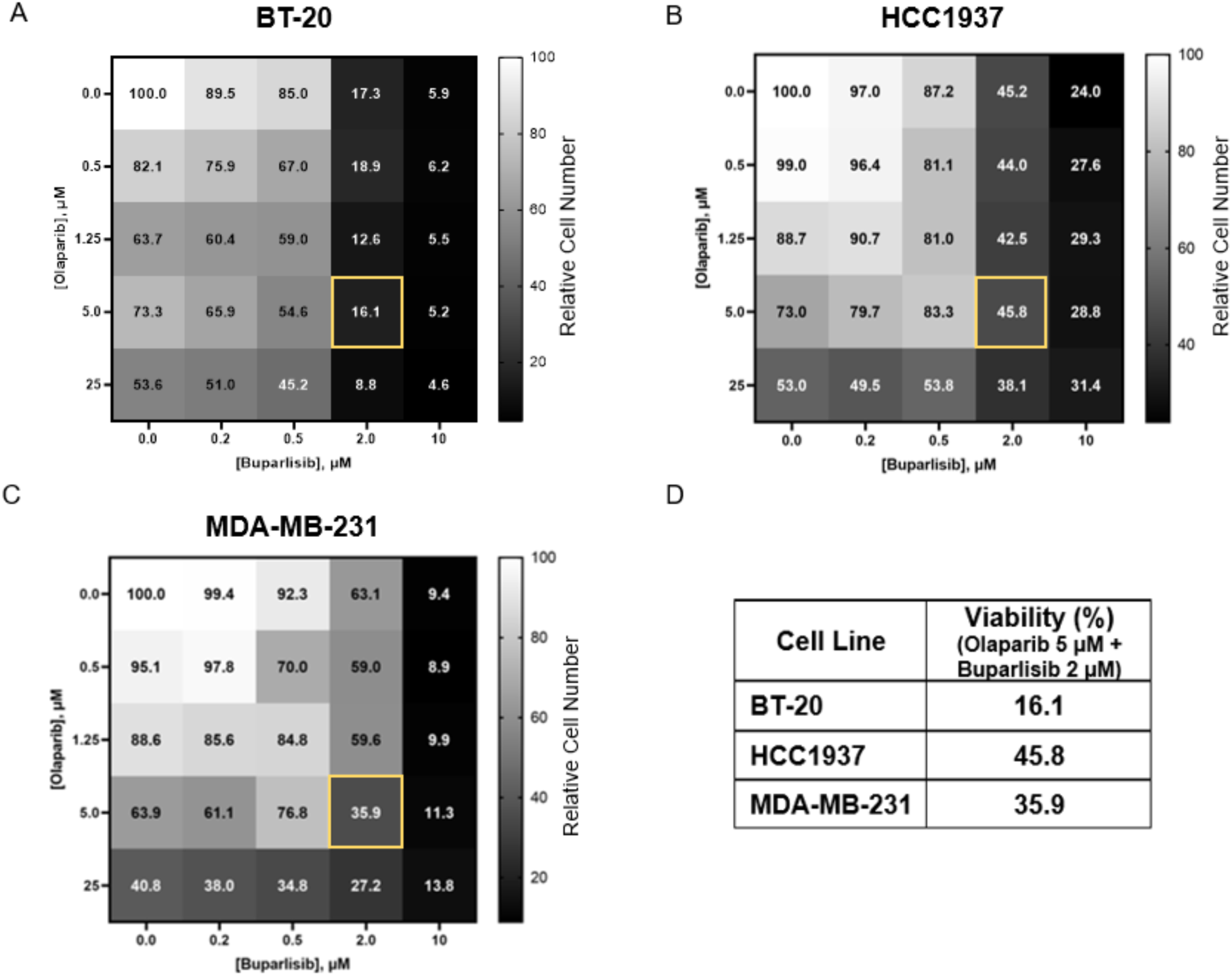
Combined PARP and PI3K inhibition synergize at low micromolar concentrations. (A – D) Viability of TNBC cell lines treated with vehicle control (VC), Olaparib (PARP inhibitor) and/or Buparlisib (PI3K inhibitor). Heatmaps indicate relative viability (color scale) with individual inhibitor treatment (top row and left-most column) and combined inhibitor treatment for BT-20 (A), HCC1937 (B) and MDA-MB-231 (C) cell lines. Concentrations used in this study are highlighted in yellow in (A – C); viability at this inhibitor combination summarized in (D). Cells were treated for 72 hrs, stained with propidium iodide and counted. Data is mean of n = 3 or more, normalized to VC.

### Initial sensitizing delivery of inhibitors elicits a cell line-specific decrease in cell viability versus simultaneous treatment

In studying the synergistic effects of combined PARPi/PI3Ki on BC cells, Juvekar et al. found that PI3K inhibition disrupts the non-oxidative pentose-phosphate pathway in BC cells, impairing nucleotide synthesis and resulting in DNA damage, replication stress, and subsequent increased PAR expression, ultimately sensitizing the cells to PARP inhibition (Juvekar et al., 2016). We asked whether sequential treatment with one drug before the other could induce a phenotypic transition in the cells which the second agent could then leverage, as has been shown in the case of taxane treatment followed by SF kinase inhibition in breast cancer (Goldman et al., 2015). To probe whether priming TNBC cells with temporally sequenced inhibition of PI3K prior to PARP inhibition, or vice versa, is more effective at inducing cell death than our simultaneous treatment approach, we exposed BT-20, HCC1937 and MDA-MB-231 cells to the following treatment conditions for a total of 72 hrs each, using the PARPi at 5 μM and the PI3Ki at 2 μM: DMSO vehicle control; individual PARP or PI3K inhibition, and combined PARP/PI3K inhibition (Olaparib/Buparlisib, or OB) (Fig. 3). For temporal sequencing, we divided the 72-hr period into 24-hr treatment periods using one inhibitor for 24 hrs followed by 48 hrs with the other, or 48 hrs first followed by 24 hrs, either replacing the initial inhibitor with the second one, or adding the second to the first at these timepoints (Fig. 3a). In all cases of temporally-staggered treatment, at the 24- or 48-hr marks, media was changed to fresh media including either the second inhibitor only, or both inhibitors as indicated. At 72 hr, cells were then fixed, stained with propidium iodide and counted using CellProfiler, and data was normalized to VC.

**Figure 3:**
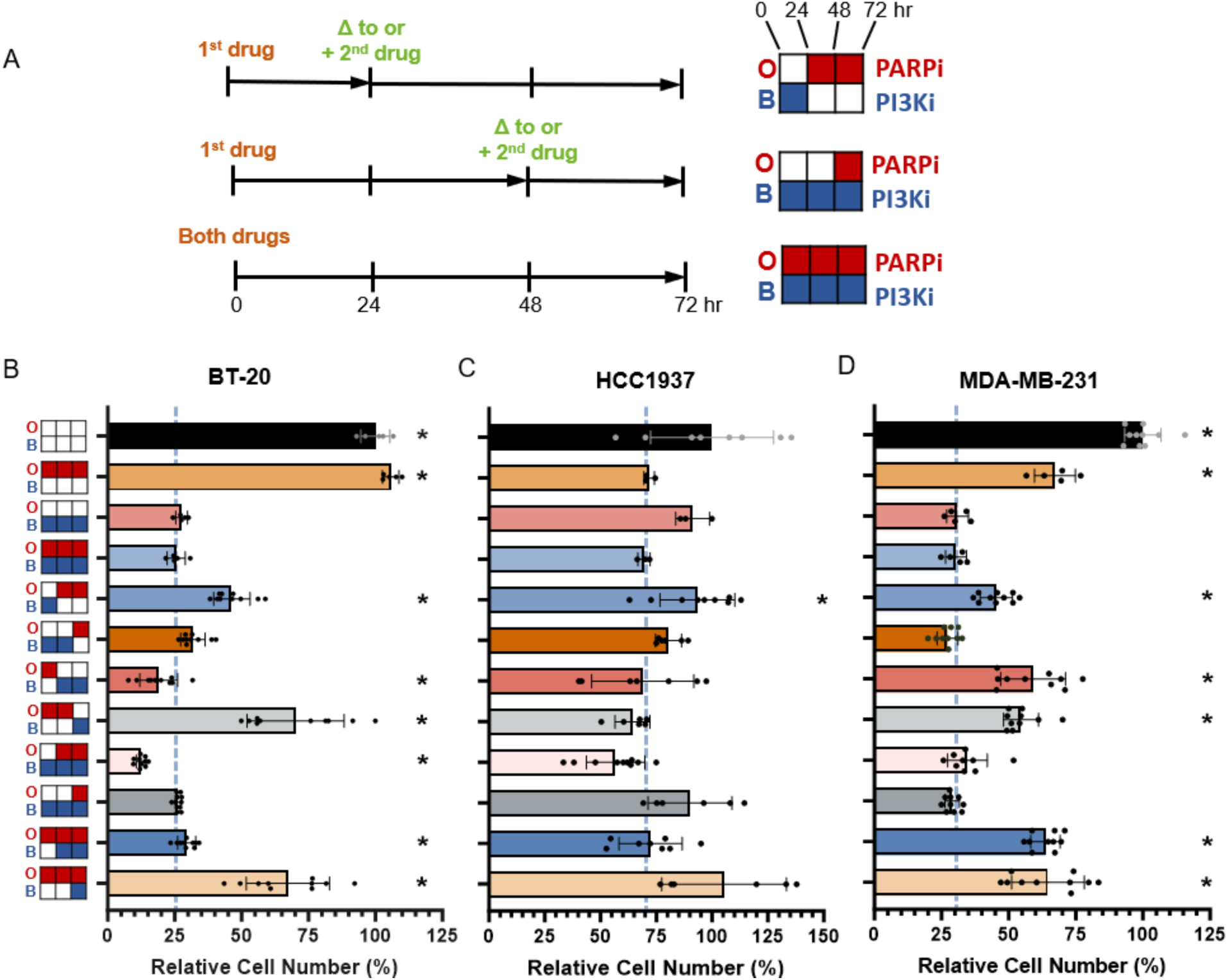
Identifying critical temporal windows of treatment in three TNBC cell lines. Scheme of treatment schedule in (A), left, with treatment key, right. Left: “+” indicates second drug was added to first; “Δ” indicates drug change. Top right: Buparlisib (BUP or B; PI3K inhibitor) 24 hrs switched to Olaparib (OLA or O; PARP inhibitor) 48 hrs; middle: BUP 72 hrs, OLA added at 48 hr; bottom: combined 72 hr treatment used in this study (OB). Cells were cultured in growth medium and treated with DMSO 0.1% vehicle control (VC) or 5 μM OLA and/or 2 μM BUP for 72 hr total. Viability studies using temporally staggered treatments for BT-20 (B), HCC1937 (C) and MDA-MB-231 (D) TNBC cell lines. Cells were fixed, stained with propidium iodide, counted and normalized to VC data (black bars). Dashed blue lines indicate 72 hr OB viability; asterisks indicate significant difference from OB. Data is mean of n = 3 or more ±SD, normalized to VC; Welch’s t-test, * p = <0.05.

We found that only two of sequential approaches significantly improved on consistent OB treatment: in BT-20 cells (Fig. 3b). OB resulted in 25.6% viability while initial PARPi treatment for 24 hrs, then switched to PI3Ki for 48 hrs, yielded 19.0% viability, and initial PI3Ki treatment for 24 hrs, with PARPi added for 48 hrs, yielded 12.5% viability (both Welch’s t-test, * p <0.05). While this unique sensitivity to temporally-sequenced inhibitors in BT-20 cells may be attributed to their pronounced vulnerability to PI3K inhibition, treating them with PI3K inhibition alone left them with slightly higher viability than OB (ns). Given this cell line is BRCA1 competent, their enhanced sensitivity to priming with PARP inhibition may be due to Buparlisib’s subsequent effects in downregulating both BRCA1 and PARP1 (Y. Li et al., 2021), the kinase’s activating mutations enhancing this effect. The 1937 cells (Fig. 3c) exhibited an overall lower sensitivity to both agents and, similar to a BT-20 trend, only PI3Ki for 24 hr, with PARPi added for 48 hr, reduced cell viability more than 10% over OB: 56.8% viability vs. 69.5% respectively (not significant). Indeed, initial PI3K inhibition may have re-established “BRCAness” in this *BRCA1*-mutated cell line (Abbotts, Dellomo, & Rassool, 2022). In MDA-MB-231 cells (Fig. 3d), we did not find a significant decrease in viability with any of our temporally sequenced conditions, although we observed the lowest viability of 27.3% with PI3K inhibition for the first 48 hr followed by PARP inhibition for 24 hr, as opposed to 30.5% viability with OB (ns). Given these cells’ proficiency in both PI3K and BRCA1, their relative insensitivity to temporal treatments could be expected. Taken together, these results suggest the efficacy of treatment is dependent upon the cells’ mutational profiles, with certain initial PI3Ki treatments providing a small advantage, indicating a PI3K-initiated depletion of nucleotide pools may sensitize TNBC cells to PARP inhibition (Juvekar et al., 2016). The BT-20 cell line, with two *PIK3CA* mutations, is exquisitely sensitive to PI3Ki, yet is the only cell line to exhibit an advantage in priming with PARPi, which may sensitize the cells to a subsequent PI3Ki-induced HR impairment mechanism identified by Ibrahim et al. (Ibrahim et al., 2012). Given these results, we elected to continue with consistent OB treatment.

### Repeat treatment with combined PARP/PI3K inhibition reveals drug-tolerant sub-populations

Drug treatment at its EC_50_ concentration for a given cell population by definition allows approximately 50% of the cells to survive. Since chemo-resistance can be defined as the intrinsic or acquired ability of cancer cells to survive treatment (Ramos, Sadeghi, & Tabatabaeian, 2021; Zhao, 2016), we drew upon our temporal drug sequencing analysis above to subject the ∼50% of cells surviving one round of 72-hr OB treatment to additional rounds. We asked whether re-treatment with combined PARP/PI3K inhibition at EC_50_ concentration would sensitize surviving TNBC cells to further drug exposure or conversely would activate intrinsic or acquired resistance mechanisms. Repeat, low-dose chemotherapy with little to no rest in between treatments, a type of metronomic therapy, is known to be effective in breast cancer (Munzone & Colleoni, 2015). To investigate, we exposed the three cell lines to OB treatment for three rounds of 72 hrs each, with 24-hr drug holidays in between each round (Fig. 4a), during which time media containing inhibitors or vehicle (DMSO) was changed to fresh growth media. At the conclusion of the third 72-hr treatment period, cells were fixed, stained with propidium iodide, imaged and counted with CellProfiler. Data was normalized to first-round VC.

**Figure 4.**
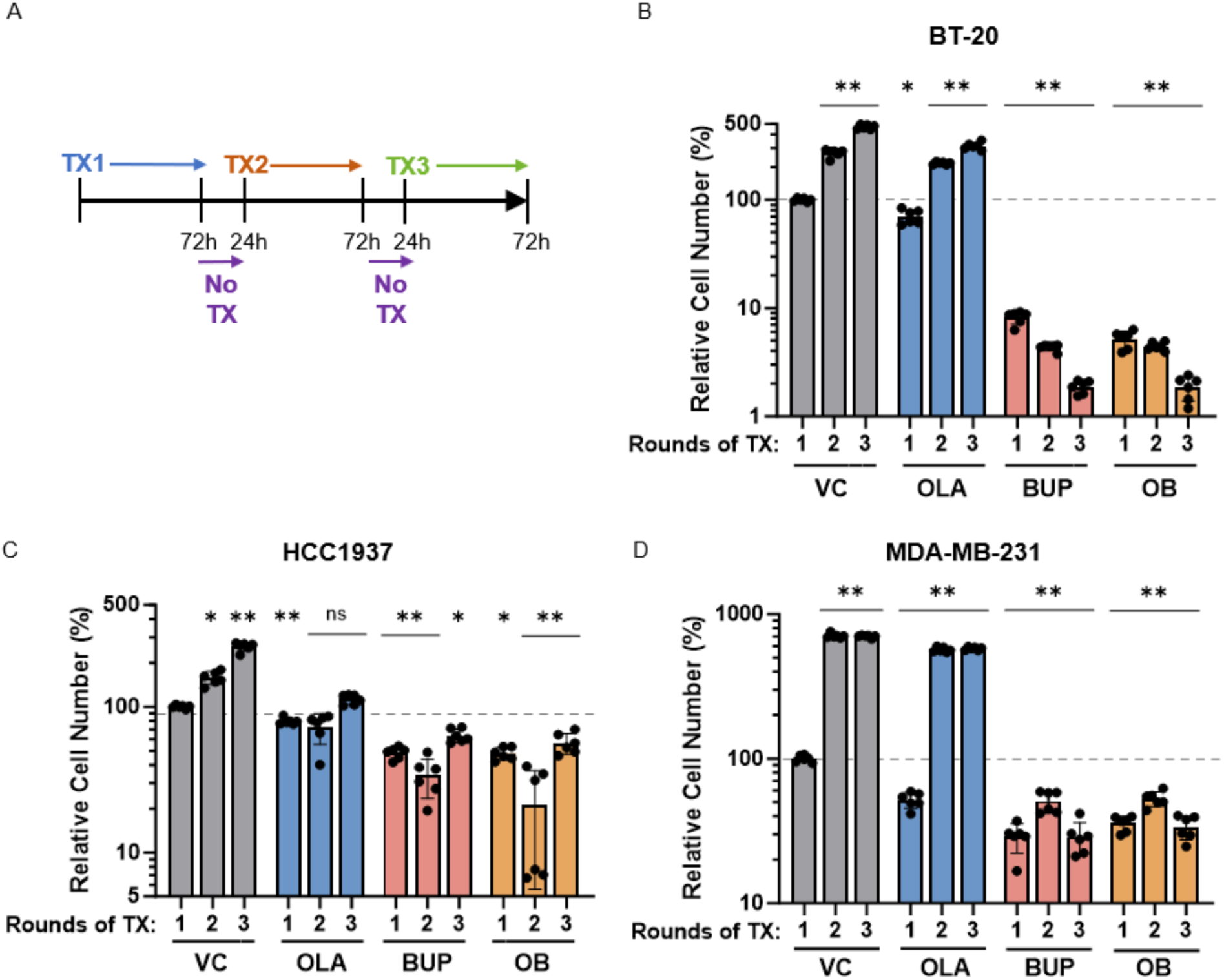
Individual and combined PARPi/PI3Ki treatment potentiation by repeated administration. Scheme of repeat 72 hr treatment schedule with 24 hr drug holidays (A); TX, treatment with vehicle control (VC) or single or combined inhibitors. Cultured cells were treated with DMSO (VC) or 5 μM Olaparib (OLA; PARP inhibitor) and/or 2 μM Buparlisib (BUP; PI3K inhibitor), or both drugs simultaneously (OB), for three consecutive courses of 72 hrs with 24 hrs drug holidays in between. Viability at each time point is shown for BT-20 (B), HCC1937 (C) and MDA-MB-231 (D) cell lines. After each 72 hr round, cells were fixed, stained with propidium iodide, counted, and data was normalized to first round VC (dashed grey line); asterisks indicate significant difference from first round VC. Data is mean of n = 6 or more ± SD; Welch’s t-test; ** p ≤ 0.0005; * p < 0.005.

In this context, BT-20 cells (Fig. 4b) were not as vulnerable to PARP inhibition as they were to PI3K inhibition and combined PARP/PI3K inhibition, which cytotoxic effects increased with each round of treatment. HCC1937 cells (Fig. 4c), which showed the greatest resistance among the three cell lines in our dual-drug studies when measured by viability (Fig. 2b), demonstrated a different overall pattern. After the first round of treatment, HCC1937 cells displayed the greatest viability of the three cell lines in each condition. However, the second round showed decreases in viability in all three conditions with a recovery after the third round. MDA-MB-231 cells (Fig. 4d), with the lowest EC_50_ concentration of Olaparib in our single-drug study, exhibited a sharp reduction in viability after one round of Olaparib treatment, then recovered robustly in the next two rounds. In 231 cells, responses to repeat PI3K inhibition and OB treatment were similar to each other, with cells recovering somewhat during the second round but reducing in number after the third. Taken together, these results indicate that PI3K and dual PARP/PI3K inhibition are generally more cytotoxic in these cells at low micromolar concentrations than PARP inhibition alone, and that repeat treatment can potentiate effects at different timepoints in the different cell lines, providing insight into advantageous dynamics of repeat chemotherapy. We posited that adaptive responses to the first round of treatment might explain these difference and sought to assess the stem-like phenotypes of surviving cells after one round of treatment.

### PARP and PI3K inhibition induce stress and survival adaptations in TNBC cell lines

To probe the basis of the restricted cell viability we observed with PARP and PI3K inhibition, we examined changes in stress-activated signaling, cleaved PARP and drug efflux pump expression by Western blot (Fig. 5a). We found that cleaved PARP expression, an early process in apoptosis, increased in all three cell lines with both PI3K and combined inhibitor treatment, and also increased in 1937 cells with PARPi treatment. While phospho-Akt expression was reduced by PI3K inhibition and OB treatment versus VC in all cell lines, there was a concomitant and possibly compensatory increase in expression of phospho-ERK 1/2 in all cell lines with PARP and PI3K inhibition, though the 1937 and 231 cells treated with OB showed a decrease in phospho-ERK1/2 expression from PI3Ki treatment alone (Fig. 5b).

**Figure 5.**
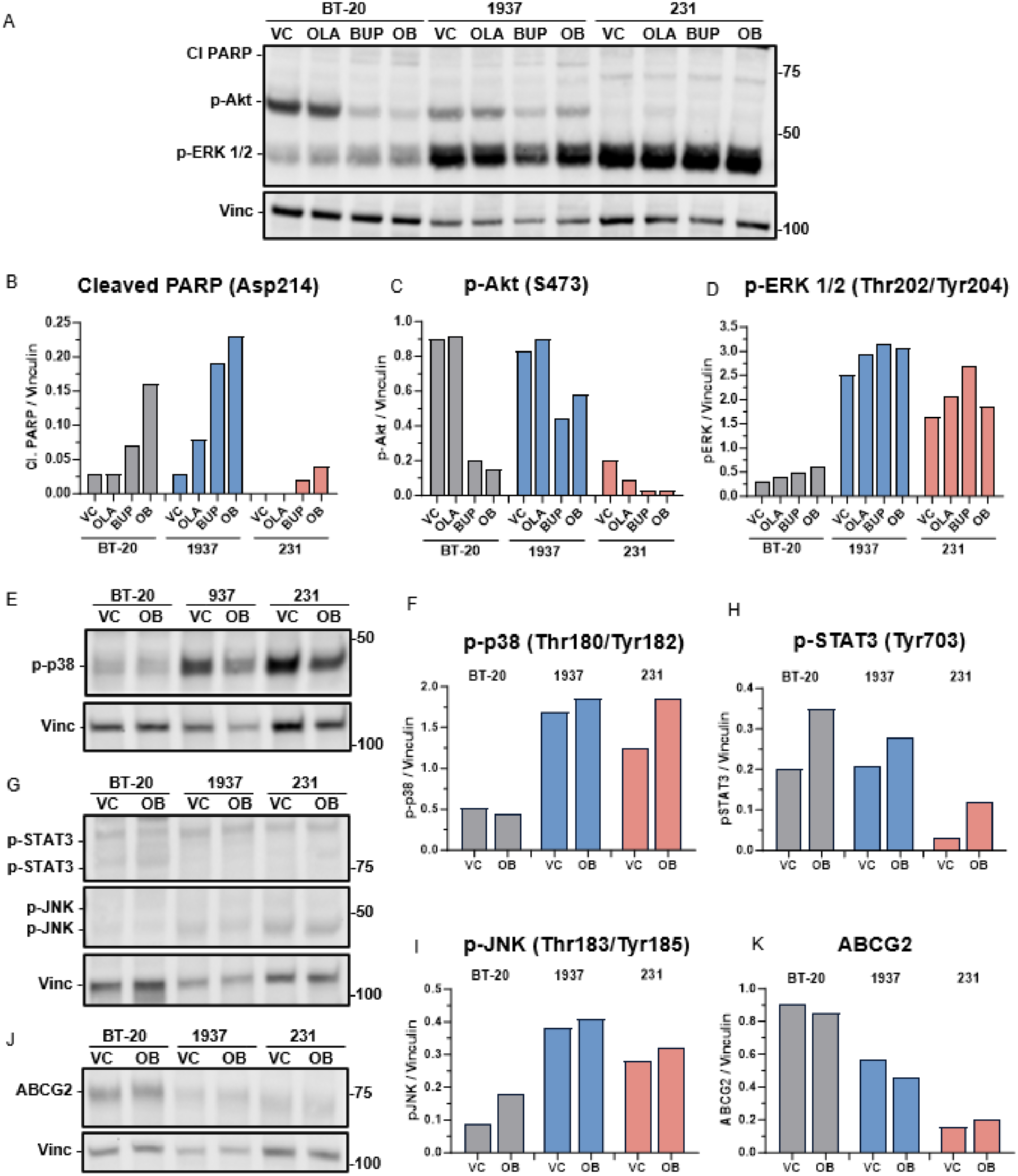
PARP and PI3K inhibition induces stress and survival signaling in TNBC cell lines with single- and dual-drug treatment. (A – D) Western blot and quantifications detailing vehicle control (VC), Olaparib (OLA; PARP inhibitor), Buparlisib (BUP; PI3K inhibitor) and combined inhibitor (OB) effects on cleaved PARP (B), phospho-Akt (S473) (C) and phospho-ERK 1/2 (Thr202/Tyr204) expression (D) in three TNBC cell lines. (E – K) Western blots and quantifications of VC and OB effects. Western blot (E) and quantification (F) of phospho-p38 (Thr180/Tyr182) expression (F). Western blot (G) and quantifications of phospho-STAT3 (Tyr703) (H) and phospho-JNK (Thr183/Tyr185) expression (I). Western blot (J) and quantification of ABCG2 expression (K). Cells were treated 72 hrs with VC, PARP and/or PI3K inhibitors; lysates probed with the indicated antibodies. Data normalized to housekeeping protein.

In testing OB treatment, we found an increase in expression of stress-related phospho-p38 MAPK in 1937 and 231 cells, and phospho-STAT3 in all three cell lines. Similarly, stress-responsive phospho-JNK increased in all three cell lines. The expression of ABCG2, an ATP binding cassette transporter protein associated with breast cancer, was reduced in BT-20 and 1937 cells with OB treatment but slightly increased in 231 cells. This may be associated with a reduction in the proportion of BCSCs with treatment, or because Buparlisib is not thought to be a substrate for ABCG2 (de Gooijer et al., 2018) while Olaparib is (Dufour et al., 2015). These results show that individual and dual inhibitor treatment induce stress- and apoptosis-related protein expression while phospho-ERK 1/2 expression is increased with both individual and dual inhibitor treatment.

### Combined PARPi/PI3Ki treatment increases proportion of CD44^+^/CD24^−/low^ and ALDH1^+^ BCSCs

Traditional chemotherapy can enrich for CD44^+^/CD24^−^ and ALDH1^+^ BCSCs (Calcagno et al., 2010) (Lee et al., 2011) (Kai et al., 2011) (Tanei et al., 2009). Since TNBC is known to be highly phenotypically heterogeneous and plastic, and harbors a CD44^+^/CD24^+^ drug-tolerant persister (DTP) subpopulation which is increased after PI3K/mTOR inhibition (Risom et al., 2018), we sought to understand how PARP and PI3K inhibitor treatment would affect these populations, and which phenotypes survive treatment. Using flow cytometry, we investigated the proportions of each population in VC and OB conditions (Fig. 6). In both the BT-20 cells (Fig. 6a,b) and 1937 cells (Fig. 6c,d), the initial CD44^+^/CD24^−^ BCSC fraction was relatively low and increased slightly with treatment: 0.5% to 1.8% and 1.2% to 4.0%, respectively (both p < 0.0001). The CD44^+^/CD24^+^ DTP subpopulation decreased slightly with treatment in both cell lines, from 99.4% to 96.7% and 98.2% to 89.7% respectively (both p < 0.0001). Conversely, the metastatic 231 cells (Fig. 6e,f), which are >99% CD44^+^/CD24 BCSCs at baseline (Zou et al., 2020), exhibited a small but significant decline in this population with treatment, from 99.3% to 98.5% and an increase in the CD44^+^/CD24^+^ DTP fraction, from 0.5% to 1.1% (both p < 0.0001). In the two cell lines which at baseline are putatively >98% DTP phenotype, this fraction was reduced with treatment while the BCSC fraction was increased. The opposite trend was observed in the metastatic 231 cells, which are predominantly BCSC at baseline: this population was reduced slightly with treatment while the DTP fraction increased. These phenotypic shifts upon treatment may underscore the adaptive plasticity of TNBC cells under chemotherapeutic stress.

**Figure 6.**
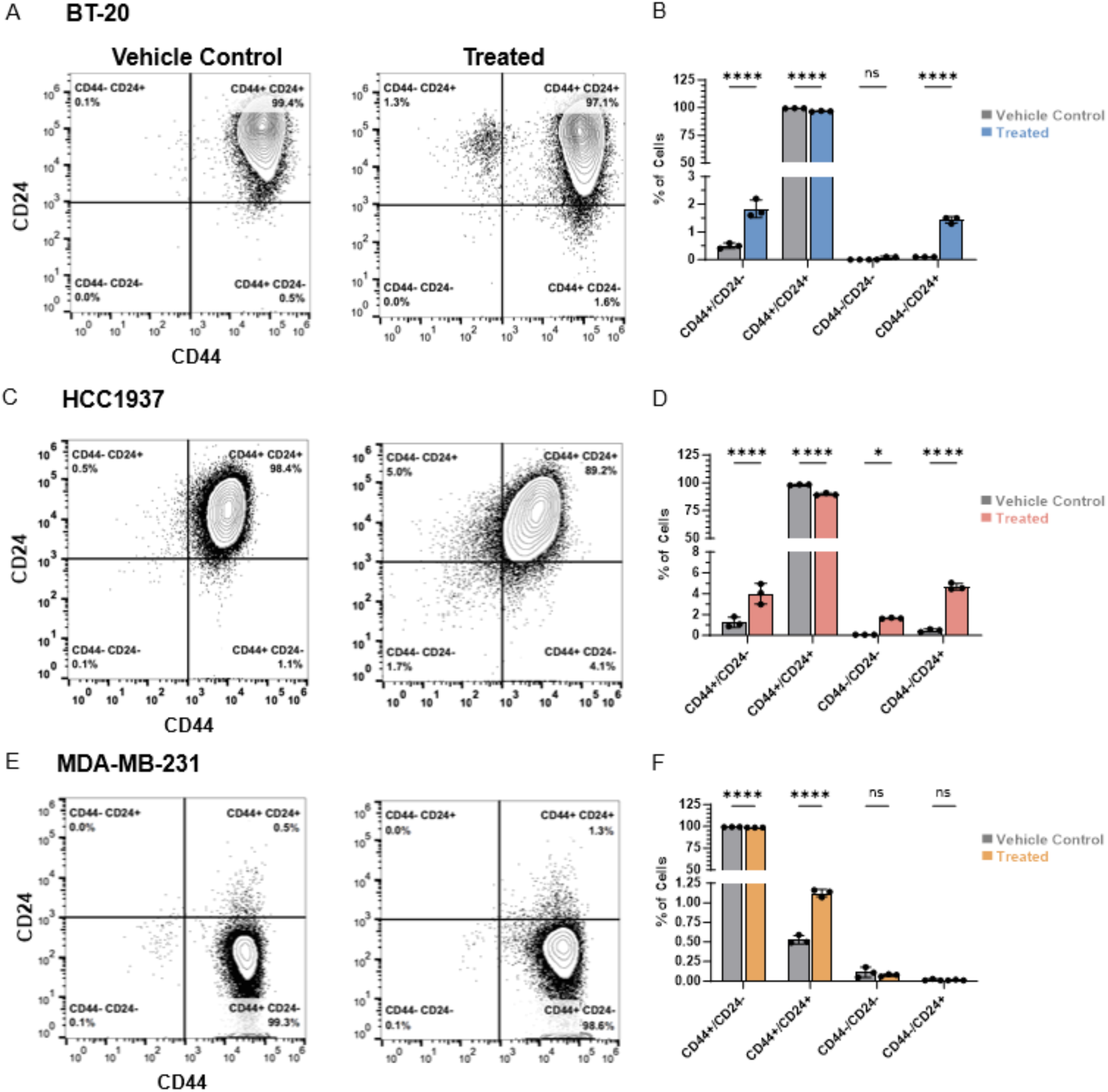
Combined PARP and PI3K inhibition induces cell line-specific increases in proportion of TNBC cancer stem cells. Flow cytometric analyses and quantifications of CSC phenotype in TNBC cells treated with vehicle control (VC) or combined PARP (Olaparib; O) and PI3K (Buparlisib; B) inhibitors (OB) for 72 hrs (A - F). Cells were dissociated and incubated with viability dye, anti-CD44 and anti-CD24 antibodies. BT-20 cell data (A – B); HCC1937 cell data (C – D); MDA-MB-231 cell data (E – F). Flow cytometry scatter plots representative of at least 3 replicates (A, C, E). Data is mean ±SD in (B, D, F). 2-way ANOVA, * p < 0.05, ** p < 0.005, *** p < 0.0005, **** p< 0.0001.

BCSCs have been shown to reversibly transit between EMT and MET states, with the CD44^+^/CD24^−^ phenotype associated with mesenchymal cells located at the tumor invasive front, and ALDH1 expression indicating proliferative epithelial cells generally located within tumors (S. Liu et al., 2014). To understand how OB treatment affects ALDH1 expression, we used flow cytometry to assess ALDEFLUOR-labelled cells. We observed a small fraction of ALDH1^+^ cells at baseline in all three cell lines of < 0.25% which increased significantly in BT-20 (Fig. 7a,b) and 1937 (Fig. 7c,d) cell lines with treatment, to 1.0% and 5.5%, respectively (both p < 0.05). The ALDH1^+^ proportion of 231 cells increased from 0.1% to 0.2% with treatment (ns) (Fig. 7e,f). These results, indicating a shift in the proportion of CD44^+^/CD24^−^ and ALDH1-expressing cells after treatment, may indicate an enrichment of drug-tolerant cells or an induction of phenotypic transitions from BCSCs to DTPs or vice versa in cell line-specific survival adaptations. As CD24 has been found to regulate EMT in breast cancer (Lim, Lee, Shim, & Shin, 2014), this change may also be indicative of EMT/MET phenotypic switching or a hybrid state in CD24^-^ cells (Bontemps et al., 2023). While it has been proposed that these types of drug-tolerant adaptations in BC are not due to Darwinian selection of more fit clones but to reversible plastic cell state transitions (Risom et al., 2018), additional longitudinal single-cell studies are necessary to illuminate the mechanisms at play.

**Figure 7.**
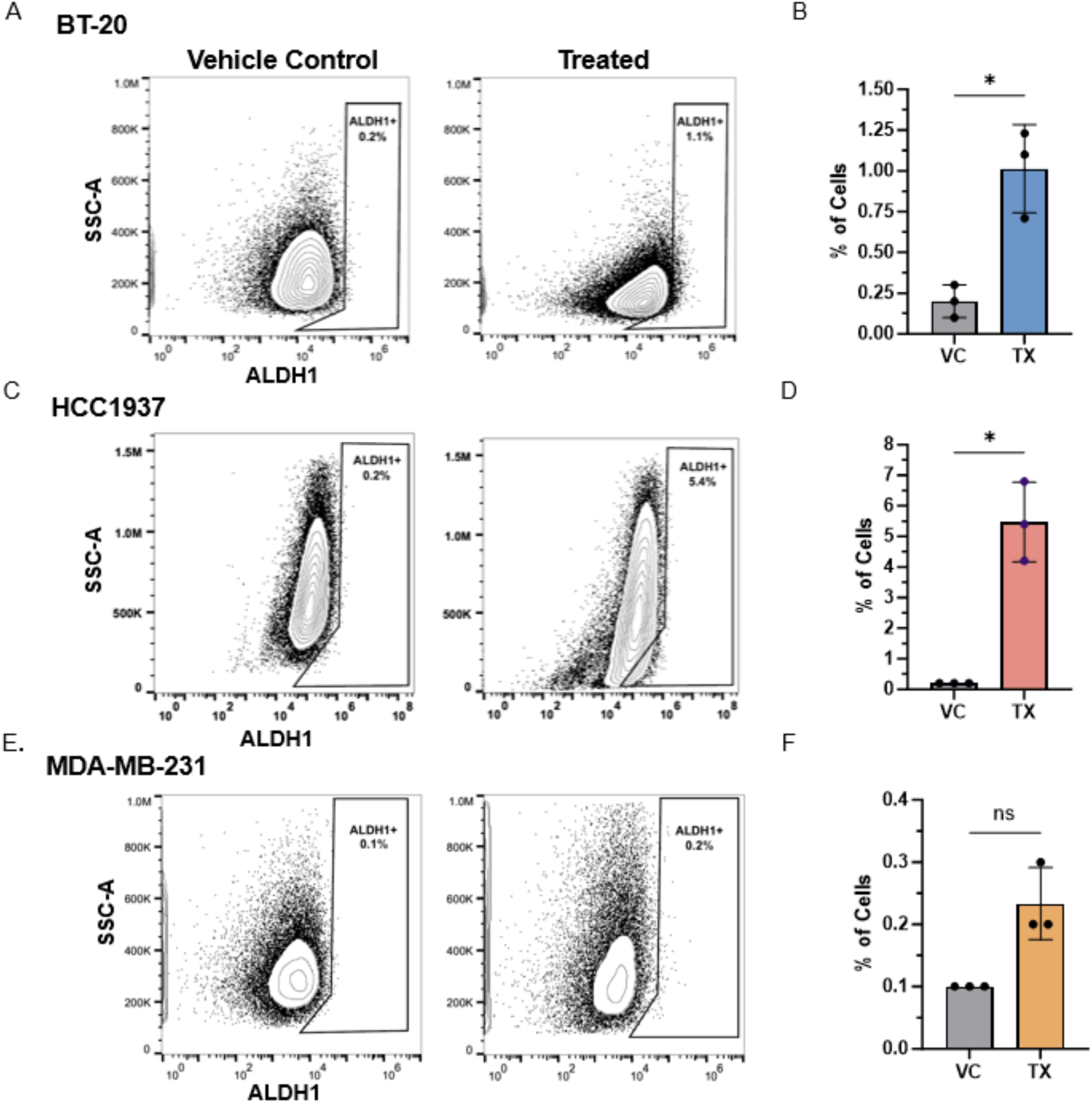
ALDH1 expression is increased with combined PARP and PI3K inhibition in two TNBC cell lines. Flow cytometric analyses and quantifications of ALDH1^+^ frequency in TNBC cells treated with vehicle control (VC) or combined PARP (Olaparib; O) and PI3K (Buparlisib; B) inhibitors (OB) for 72 hrs (A - F). BT-20 cell data (A - B); HCC1937 cell data (C – D); MDA-MB-231 cell data (E – F). Cells were dissociated and incubated with viability dye and the ALDH1 marker, ALDEFLUOR. Flow cytometry scatter plots representative of at least 3 replicates (A, C, E). Quantifications of ALDH1^+^ cell frequency (B, D, F); data is mean ±SD. Welch’s t-test, * p < 0.05.

### DNA transcription is restricted with PARPi/PI3Ki treatment

To further explore the mechanisms by which combined PARP/PI3K inhibitor treatment restricts cell viability, we examined BrdU incorporation by flow cytometry in VC and treated cells (Fig. 8). Increases in the proportion of sub-G1 cells with treatment were significant in 1937 and 231cells (Figs. 8c,d and 8e,f, respectively), (both p < 0.005) as were reductions in the S-phase fractions (both p < 0.0001). All three cell lines increased their proportions of G2-phase cells with treatment (p < 0.0005 in 231 cells). Arrest in G2 is correlated with unresolved DNA damage (DiPaola, 2002) and may indicate delay or abrogation of entry into mitosis. With mutated *CDKN2A*, BT-20 and 231 cells may maintain aberrant cell cycle dynamics and reduced apoptotic responses to treatment. These results indicate that DNA transcription in all cell lines was reduced with combined inhibitor treatment, accompanied by increases in G2 and sub-G1 populations, with BT-20 cells registering the least disruption in cell cycle dynamics of the cell lines after treatment. Ki67 flow cytometry assays provided confirmation in one cell type of reduced S-phase dynamics and an increase in apoptotic cells (Fig. S2).

**Figure 8.**
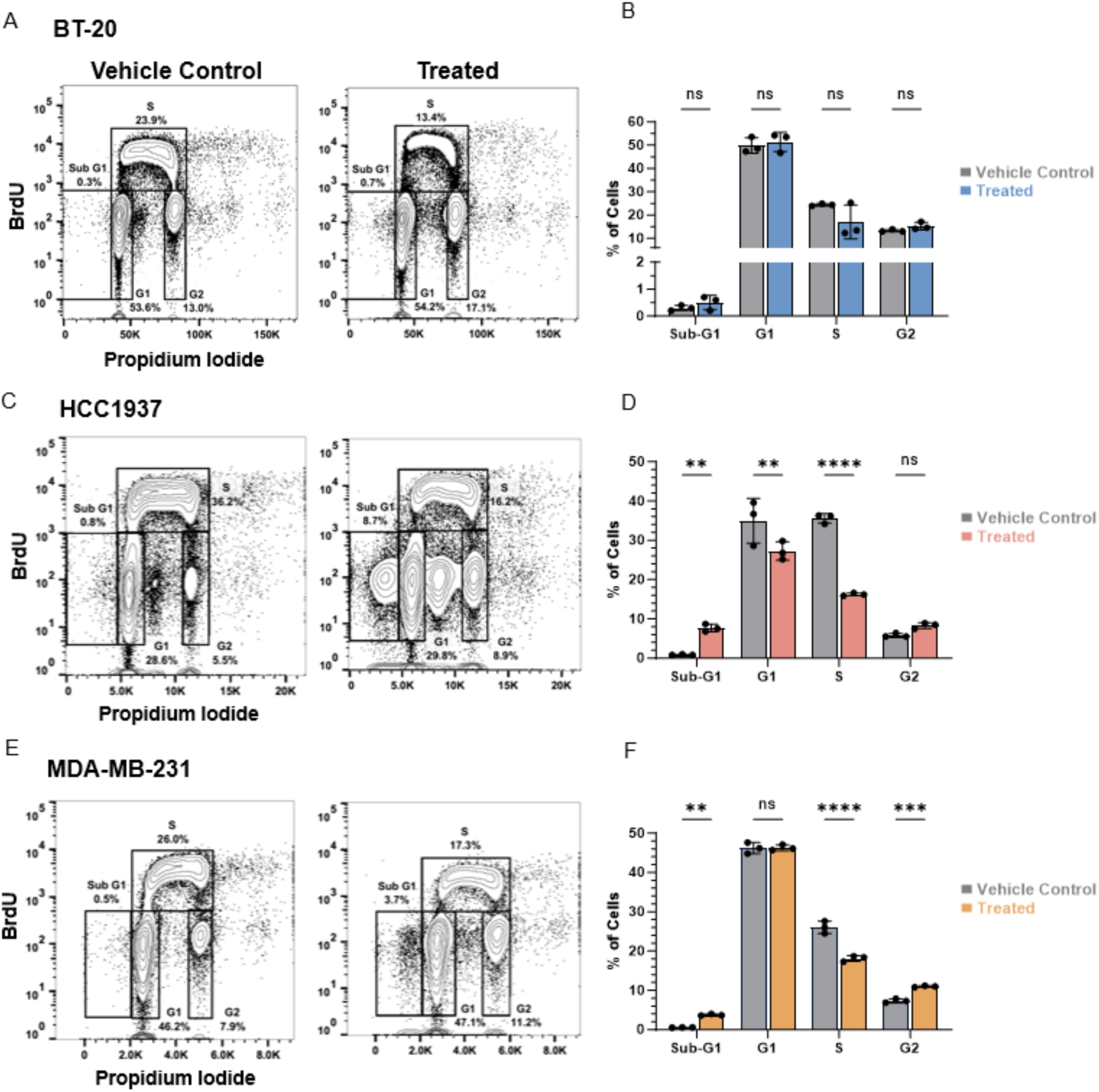
DNA synthesis in two TNBC cell lines is reduced by combined PARP and PI3K inhibition. Flow cytometric analyses and quantifications of BrdU incorporation in TNBC cells treated with combined PARP (Olaparib; O) and PI3K (Buparlisib; B) inhibitors (OB) for 72 hrs (A - F). Cells were dissociated then incubated with BrdU; stained with anti-BrdU antibody and propidium iodide. BT-20 cell data (A - B); HCC1937 cell data (C – D); MDA-MB-231 cell data (E – F). Flow cytometry scatter plots representative of at least 3 replicates (A, C, E). Quantifications of cell frequency detected in each cell cycle stage in (B, D, F); data is mean ±SD. 2-way ANOVA, * p < 0.05, ** p < 0.005, *** p < 0.0005, **** p< 0.0001.

### Proportions of live and early apoptotic cells are significantly altered in three TNBC cell lines

We examined the effects of dual inhibitor treatment on apoptosis using Annexin V stain and flow cytometry (Fig. 9). In all three cell lines the proportion of cells undergoing early apoptosis increased significantly and the proportion of live cells decreased significantly with OB treatment. However, the 1937 cell line, with the highest percentage of viable cells in our dual drug tests of 45.8% (Fig. 2b), and overall lowest drug sensitivity in our temporal treatment and re-treatment models (Figs. 3c and 4c respectively), showed the smallest decrease in the proportion of live cells and smallest increases in proportions of dead and early and late apoptotic cells of the three cell lines (Fig. 9c,d). In contrast, BT-20 and 231 cells displayed dramatic decreases in live cells and increases in early apoptotic cells (all p < 0.0001). The 231 cells, which are ∼99% BCSCs at baseline, did not exhibit enhanced resistance to apoptosis relative to the other cell lines, and indeed exhibited the greatest increase in the proportion of dead cells with treatment, increasing from 3.0% to 16.1% (p < 0.0001). In our observations, our 72-hr treatment endpoint captured apoptotic processes at varying stages for each of the three cell lines and may have captured recovering populations as well. Mutational profiles and diverse responses to combined treatment may impact apoptotic timing and severity.

**Figure 9.**
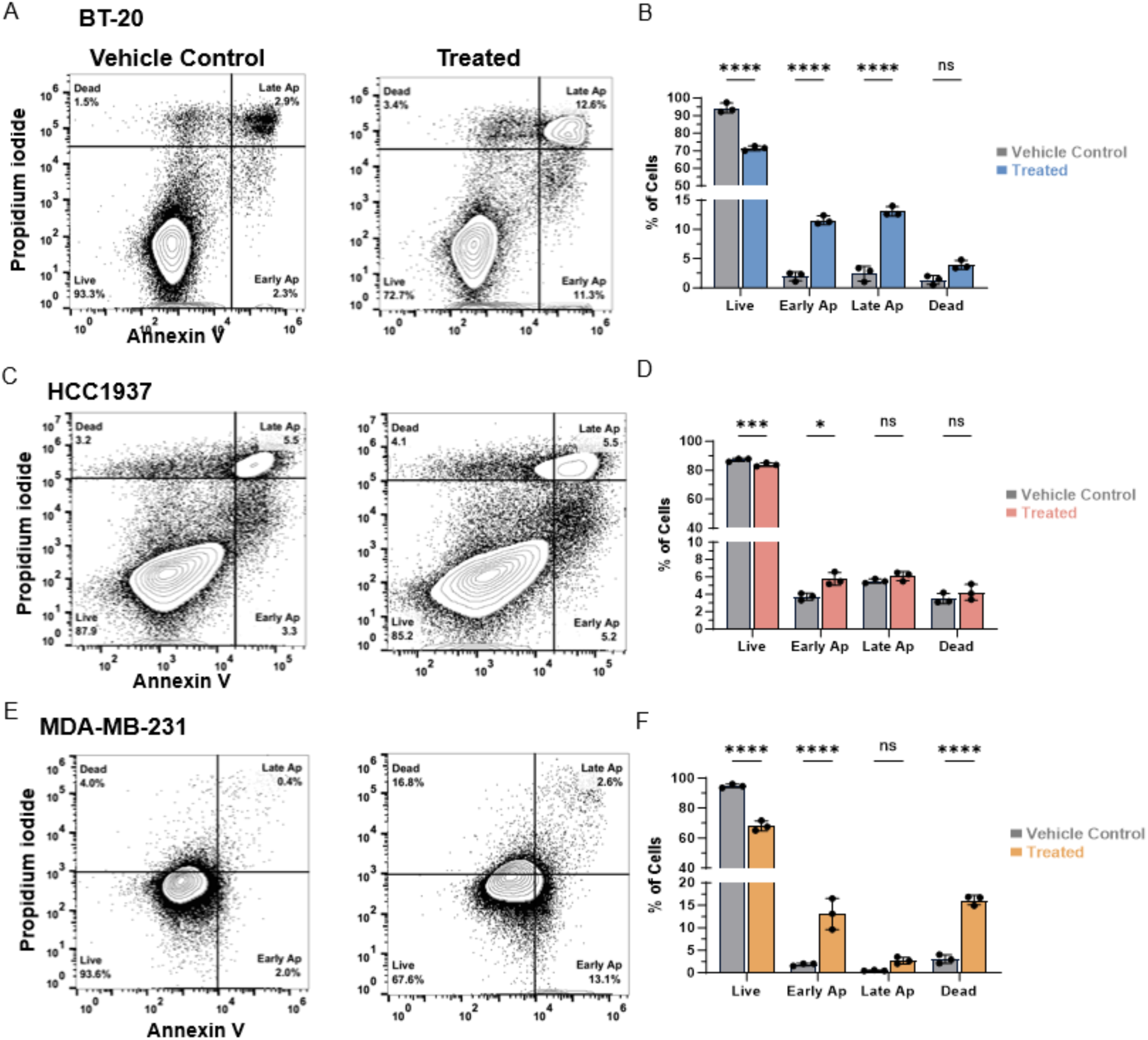
Combined PARP and PI3K inhibition reduces cell viability and induces apoptosis in three TNBC cell lines. Flow cytometric analyses and quantifications of apoptotic TNBC cells treated with vehicle control (VC) or combined PARP (Olaparib; O) and PI3K (Buparlisib; B) inhibitors (OB) for 72 hrs (A – F). BT-20 cell data (A – B); HCC1937 cell data (C – D); MDA-MB-231 cell data (E – F). Cells were dissociated, dead cells retained, and incubated with Annexin V and propidium iodide. Flow cytometry scatter plots representative of at least 3 replicates (A, C, E). Data Is mean ±SD in (B, D, F). 2-way ANOVA; * p < 0.05 ** p < 0.005; *** p < 0.0005; **** p < 0.0001.

## Discussion

Here we evaluated therapeutic vulnerabilities in TNBC stem-like cells distinct from those targeted by standard-of-care chemotherapies. Previous work has revealed that chemotherapy can trigger an increase in the proportion of CD44^+^/CD24^−^ and ALDH1 expressing BCSCs (Calcagno et al., 2010; Creighton et al., 2009; Ghanbari et al., 2016; Jaggupilli et al., 2022; Lee et al., 2011). Our use of combined PARP and PI3K inhibition (Olaparib and Buparlisib, respectively) to target BCSCs is a novel approach, building on the work of Juvekar, et al. (Juvekar et al., 2012; Juvekar et al., 2016) in which this combination is shown to inhibit BRCA1-mutated BC cells and to synergize to deplete nucleoside pools, as well as the findings of Zhou, et al., which implicate PI3K inhibition in the reduction of CSCs in TNBC (H. Zhou et al., 2019).

We found that combined PARP/Pi3K inhibition decreases cell viability, restricts DNA transcription, induces stress and survival adaptations, and after one round of treatment maintains or increases the proportion of CD44^+^/CD24^−/low^ and ALDH1^+^ CSCs in a small but significant manner. We also found that sequential drug delivery and re-treatment protocols decreased cell viability in certain conditions across cell lines. This increase in CSCs after treatment comports with previous work using conventional chemotherapies, and expands our understanding of the roles of PARP and PI3K in BCSC chemoresistance (Calcagno et al., 2010; Creighton et al., 2009; Ghanbari, Azadbakht, Vesi-Raygani, & Khazaei, 2016; Jaggupilli et al., 2022; Lee et al., 2011). Identifying CSC-associated cell surface markers and metabolic signatures, and designing appropriate targeted therapies are currently areas of intense investigation (Ali et al., 2024).

Indeed, as knockdown of CD44 in BC results in a loss of stemness and induces sensitivity to chemotherapy (Pham et al., 2011), the inhibition of this molecule or its effectors is of interest in chemotherapeutic development (Girda et al., 2022; M. Li et al., 2021; Xu, Niu, Yuan, Wu, & Liu, 2020). Interestingly, overexpression of CD24 in TNBC can enhance chemo-sensitivity (Deng et al., 2017), and TNBC is known to encompass a CD44^+^/CD24^+^ drug-tolerant persister (DTP) subpopulation (Risom et al., 2018), which we found to increase in two of our cell lines with treatment. Because high CD24 expression is associated with poor prognosis in TNBC (Kwon et al., 2015), it is a poor target for therapy evaluation. As of yet, the safety and efficacy of ALDH inhibitors has not been established (Wei et al., 2022).

Olaparib has been FDA-approved for the treatment of germline *BRCA*-mutated (g*BRCA*m), HER2^−^ BC and g*BRCA*m ovarian cancer (Mandapati & Lukong, 2023; Singh, Parveen, & Yadav, 2021), and Buparlisib and similar PI3K inhibitors are in clinical trials for use in BC and ovarian cancer, with Alpelisib approved for use in *PIK3CA*-mutated BCs. (Yu et al., 2023) As ovarian cancers are also linked to germline and somatic *BRCA1* mutations (Berchuck et al., 1998; Stratton et al., 1997), and are thought to harbor CSCs like BC (Cioffi et al., 2015; Krolewska-Daszczynska, Wendlocha, Smycz-Kubanska, Stepien, & Mielczarek-Palacz, 2022), this combination of drugs may prove effective beyond the BC context. Following this logic, combined Olaparib and Alpelisib has shown promise in a Phase 1b clinical trial in breast and ovarian cancers (Konstantinopoulos et al., 2019).

It will be advantageous to test our approach in patient samples and in PDX mouse models, and longitudinal studies of treated cells would improve our understanding of adaptations to treatment. Given the complex interactions of PARP and PI3K proteins, much work is needed to elucidate their cognate ligands and convergent signaling pathways in TNBC. Our results have therapeutic implications due to our design of effective treatment protocols using combined PARP and PI3K inhibition, and given the additive effect we identified, this combination may show efficacy at lower clinical doses with the potential to create a safer therapeutic profile.

## Methods

### Common Reagents

All common laboratory reagents were obtained from VWR (Radnor, PA), Sigma-Aldrich (Burlington, MA) or ThermoFisher (Carlsbad, CA).

### Cell Culture

All cells were maintained in two-dimensional culture at 5 x 10^4^ cells/mL and 37°C in 5% CO_2_ in a humidified chamber. BT-20 cells were cultured in growth medium using Minimum Essential Medium (MEM; Corning, Inc., Oneonta, NY, #10-009-CV) with 10% fetal bovine serum (FBS; Corning, Inc., #35-010-CV) and 1% penicillin/streptomycin (ThermoFisher, #15140122). Growth medium for both HCC1937 and MDA-MB-231 cells was Roswell Park Memorial Institute 1640 (RPMI 1640; Corning, Inc., #10-041-CVR) with 10% FBS and 1% penicillin/streptomycin. Cancer cell lines were passaged fewer than 20 times. MDA-MB-231 cells were a generous gift from Claudia Fischbach-Teschl of Cornell University. Other cell lines were purchased from American Type Culture Collection (atcc.org, Manassas, VA).

### Cell Viability

Olaparib (PARPi) and Buparlisib (PI3Ki) were obtained from Selleck Chemicals LLC (Houston, TX, #S1060 and #S2247 respectively) and reconstituted in DMSO (Sigma-Aldrich, #D8418). Cells were plated at a density of 5 x 10^4^ cells/mL and allowed to attach for 24 hrs, then treated with Olaparib and/or Buparlisib at the indicated concentrations up to 72 hrs. Vehicle control cells were maintained in normal growth medium supplemented with 0.1% DMSO, and DMSO was added to inhibitor-treated cells to normalize its concentration. Cells were not included in in 96-well plate edge wells to minimize edge effects. Methanol fixation was followed by propidium iodide staining (ThermoFisher, #P3566), cells were counted using CellProfiler (Broad Institute, Cambridge, MA) and data was normalized to vehicle control cells. MATLAB (MathWorks, Natick, MA) and Prism (GraphPad Software, Boston, MA) were used for heatmap graphs, and Prism and Excel were used for statistical analyses.

### Imaging

Nikon Eclipse Ti-E Microscope (MVI, Avon, MA) with Spectra X or Celesta light sources was used to capture digital images. For viability studies, propidium iodide was used to stain nuclei; images were captured in the TRITC channel; thresholding performed in native NIS Elements software; nuclear segmentation and automated counting were performed using CellProfiler.

### Quantitative Immunoblotting

Cells were lysed with RIPA or NP-40 buffer as appropriate for cellular compartments of target proteins. RIPA: 50 mM Tris-HCl, 150 mM NaCl, 1% Triton X-100, 0.5% sodium deoxycholate, 0.1% SDS, 5 mM EDTA supplemented with 10 μg/mL aprotinin, 10 μg/mL leupeptin, 1 μg/mL pepstatin, 1 mM PMSF, 1 μg/mL microcystin-LR, 200 μM sodium orthovanadate. NP-40: 50 mM Tris-HCl, 150 mM NaCl, 0.5% NP-40 substitute, 5 mM EDTA supplemented as above. Cells were placed on ice and washed with cold PBS. Cold lysis buffer was added to each well and allowed to incubate on ice for 15 minutes. Cells were scraped and lysates were collected, then centrifuged at 4° C for 10 minutes at 15,000 x g. The supernatant was collected and protein concentration was quantified with a micro bicinchoninic acid assay kit (ThemoFisher, Waltham, MA, #23235). Lysates were electrophoresed and transferred using the ThermoFisher BOLT Bis-Tris system (#NW0008C). Each lysate was loaded with Bolt sample buffer and reducing agent onto electrophoresis gels at 10 - 20 μg of protein per lane, with chemiluminescent protein standards (Bio-Rad, Hercules, CA, #1610374 and #1610376) per manufacturer’s recommendations. Gels were run using Bolt MES running buffer at 200 V for 35 minutes. Proteins were wet-transferred at RT to a methanol-activated PVDF membrane using BOLT transfer buffer, methanol and anti-oxidant at 20 V for 60 minutes. Membranes were blocked in 5% bovine serum albumin (BSA) (Rockland Immunochemicals, #BSA-50) in Tris-buffered saline with Tween (TBST) for 1 hr with gentle rocking at RT. Primary antibodies were diluted in 5% BSA/TBST at the following concentrations: GAPDH (36 kDa, rabbit polyclonal Ab, Invitrogen #PA1-987), 1:10,000; all following are Cell Signaling Technology (Danvers, MA): ABCG2 (65–80 kDa, clone D5V2K, rabbit mAb, #42078), 1:1000; Cleaved PARP (89 kDa, Asp214, rabbit mAb, #9541), 1:1000; Phospho-Akt (Ser473, 60 kDa, clone D9E, rabbit mAb, #4060), 1:2000; Phospho-p38 (Thr180/Tyr192, 43 kDa, clone D3F9, rabbit mAb, #4511), 1:1000; Phospho-p44/42 MAPK (ERK 1/2) (Thr202/Tyr204, 42, 44 kDa, rabbit mAb, clone D13.14.4E), 1:2000; Phospho-SAPK/JNK (Thr183/Tyr185, 46 and 54 kDa, rabbit mAb, clone 81E11, #4668), 1:1000; Phospho-STAT3 (Tyr705, 79 and 86 kDa, rabbit mAb, clone D3A7, #9145), 1:2000; Vinculin (124 kDa, rabbit mAb, clone E1E9V, #13901), 1:1000. Blots were incubated with primary antibodies with gentle rocking at 4° C overnight, then washed with TBST three times for five minutes each. Blots were then incubated with goat anti-rabbit secondary antibody HRP conjugate (Jackson ImmunoResearch Laboratories, West Grove, PA, #111-005-045, 1:5,000 dilution) and HRP-Streptactin Conjugate (Bio-Rad, #1610380) at 1:10,000 dilution in TBST for 1 hr at RT, then washed again as above. Blots were then incubated in ECL substrate (Bio-Rad, #1705060) for 1 minute at RT and imaged on the ChemiDoc system (Bio-Rad, #17001401). Protein expression was normalized to housekeeping genes and quantified using ImageJ, Excel and GraphPad Prism software.

### Flow cytometry

Briefly, for CD44 and CD24 assays, cells were harvested with Accutase (Innovative Cell Technologies, Inc., San Diego, CA, #AT104) to preserve surface proteins, then treated with FC block (ThermoFisher, #14-9191-73) before staining with BioLegend (San Diego, CA) antibodies as follows: anti-CD44 antibody variously conjugated to AF700 (clone C44Mab-5, #397522), APC-Cy7 (clone IM7, #103028), FITC (clone C44Mab-5, #397518) or PE-Cy5 (clone IM7, #103010), and anti-CD24/PE (clone W20001B, #382610). DAPI (ThermoFisher, #D3571), fixable viability dye (ThermoFisher, #L34963) or propidium iodide (ThermoFisher # P3566) were used for live/dead cell discrimination as appropriate for experimental exigencies.

For ALDH1 assays, the ALDEFLUOR kit was used from STEMCELL Technologies, Inc. (Vancouver, BC, Canada, #1702), per the manufacturer’s protocol. Cells were also stained with anti-CD44 and anti-CD24 antibodies as above. For BrdU incorporation, briefly, cells were pulse-labeled with BrdU for 2 hr then stained with fixable viability dye. Cells were fixed in ice-cold 70% ethanol, then DNA was denatured with 2 N hydrochloric acid with 0.5% Triton-X in phosphate-buffered saline (Corning, # 46-013-CM). Neutralization with sodium tetraborate was followed by incubation with anti-BrdU/FITC antibody (ThermoFisher, clone BU20A, #11-5071-42). Cells were stained with propidium iodide and treated with RNase A (VWR, # VWRVE866-5ML). For Annexin V staining, briefly, cells were washed and resuspended in Annexin binding buffer (ThermoFisher, #V13246), then stained with Annexin V/FITC (ThermoFisher, # A13199) and incubated at RT. Propidium iodide was added to the cells prior to analysis.

For Ki67 assays, cells were harvested with Accutase then incubated with anti-Ki67/FITC antibody (ThermoFisher, mouse mAb, clone 7B11, #MHKI6701) and stained with propidium iodide.

Flow cytometry was performed on equipment from Cornell University’s Biotechnology core, using BD Aria Fusion, BD Melody (both Becton Dickinson and Company, Franklin Lakes, NJ), Sony MA900 (Sony Biotechnology, San Jose, CA), and Thermo Attune NxT (ThermoFisher Scientific) cytometers. Flow data analysis performed using FlowJo software (Becton Dickinson and Company).

### Statistical analyses

Unless stated otherwise, experimental data are calculated as mean values normalized to vehicle control data, with error bars representing ±SD for a minimum of three biological replicates. Differences in means between experimental groups was evaluated using unpaired or Welch’s t-tests, Brown-Forsythe and Welch’s 2-way ANOVA as appropriate, and a p-value of less than 0.05 was considered significant. GraphPad Prism, MATLAB and Microsoft Excel were used for data analysis and visualization.

## Acknowledgments

The authors would like to thank Joseph Druso, of the Fischbach-Teschl lab for assistance in culturing MDA-MB-231 cells; Lu Ling of the Fischbach-Teschl lab for assistance in culturing MCF-10A cells; and Marc Antonyak of the Cerione lab for assistance in Western blotting. We extend our thanks to Lydia Tesfa, Jaclyn Mahoney and Michael Sledziona at the Flow Cytometry Facility (RRID:SCR_021740) of the Cornell Biotechnology Resource Center for their help in performing flow cytometry experiments. This work was supported by the US National Institutes of Health (NIH) grants U54CA210184, R01CA238745, R01CA260115 and R01CA248524 (B.D.C.) and Cornell Stem Cell Training Fellowship (P.E.P.). The content is solely the responsibility of the authors and does not necessarily represent the official views of the NIH.

---

**Figure S1.**
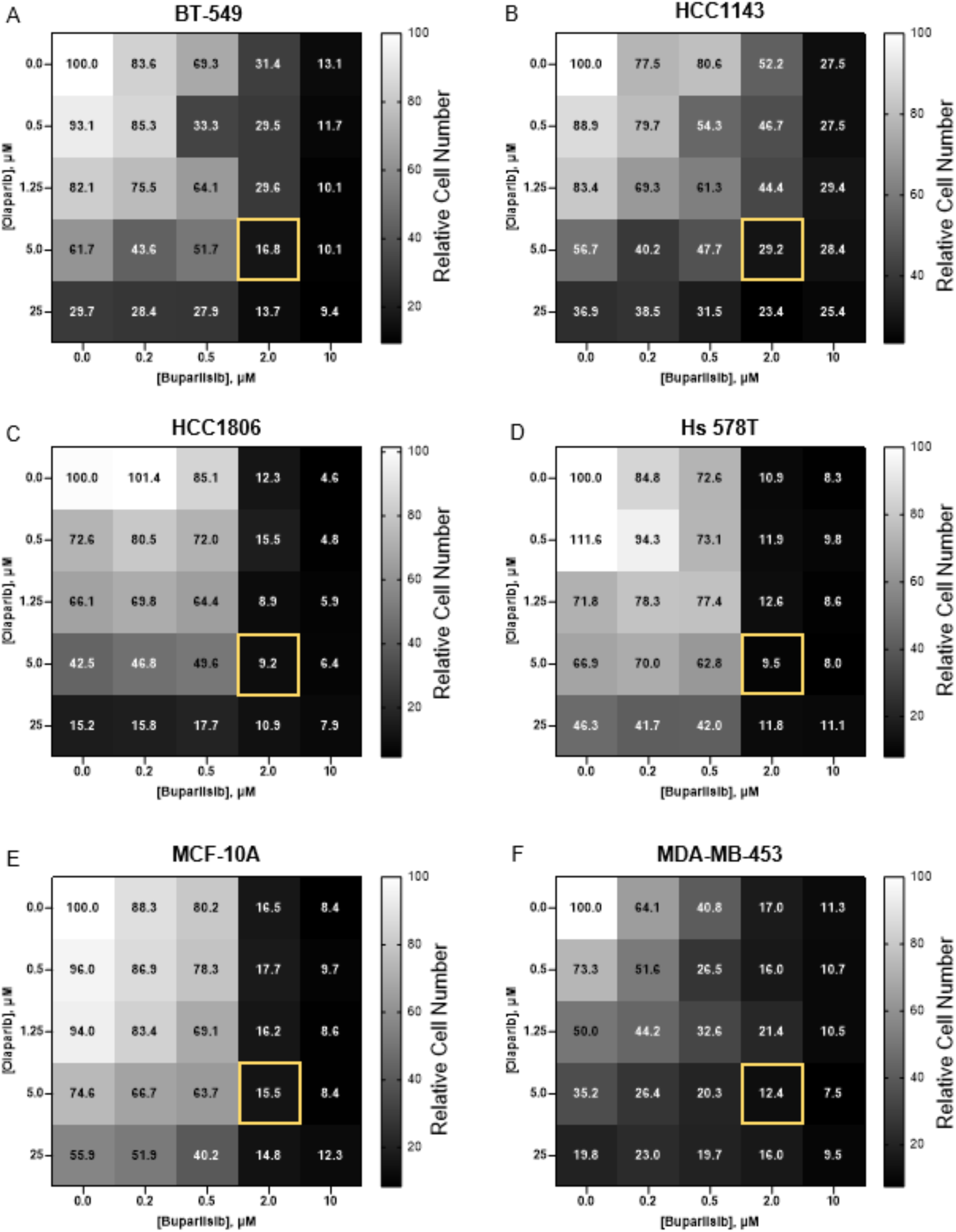
Viability heatmaps for five additional TNBC cell lines, and the non-transformed mammary epithelial cell line MCF-10A, which were evaluated for this study (A – F). Cells were treated 72 hrs with vehicle control (VC) Olaparib (PARP inhibitor) and/or Buparlisib (PI3K inhibitor). Heatmaps indicate relative viability (color scale) with individual inhibitor treatment (top row and left-most column) and combined inhibitor treatment for BT-549 (A), HCC1143 (B), HCC1806 (C), Hs 578T (D), MCF-10A (E), and MDA-MB-453 (F) cell lines. Concentrations used in this study are highlighted in yellow in (A – F). At 72 hrs cells were stained with propidium iodide and counted. Data is mean of n = 3 or more; data normalized to VC.

**Figure S2.**
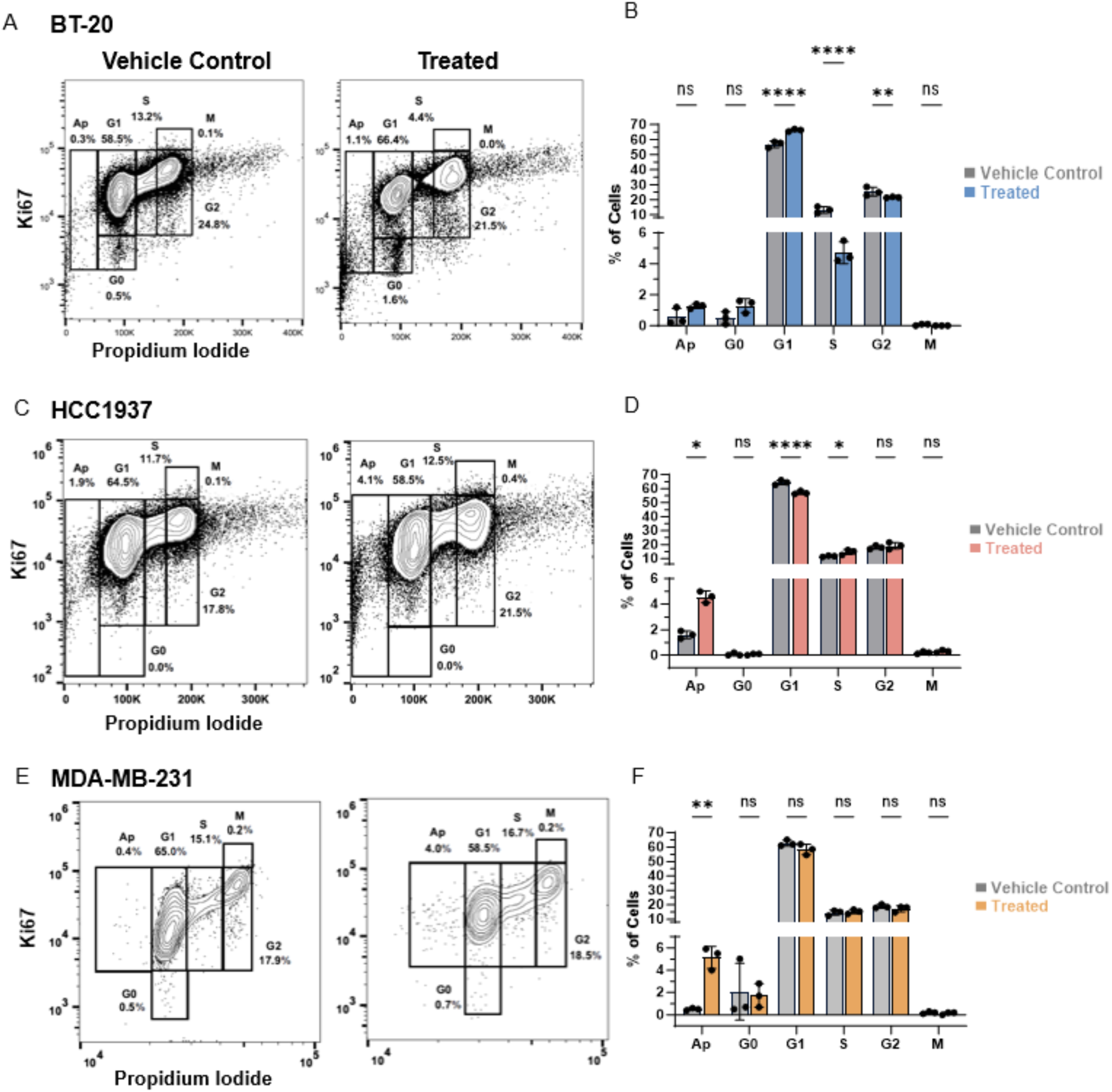
Combined PARP and PI3K inhibition induces apoptosis and decreases S-phase fraction in cell line-dependent manner. Flow cytometric analyses and quantifications of cell cycle stages in TNBC cells treated with vehicle control (VC) or combined PARP (Olaparib; O) and PI3K (Buparlisib; B) inhibitors (OB) for 72 hrs (A - F). BT-20 cell data (A - B); HCC1937 cell data (C – D); MDA-MB-231 cell data (E – F). Cells were dissociated and incubated with anti-Ki67 antibody and propidium iodide. Flow cytometry scatter plots representative of at least 3 replicates in (A, C, E). Quantifications of cells in each cell cycle stage in (B, D, F); data is mean ±SD; 2-way ANOVA, * p < 0.05, ** p < 0.005, *** p < 0.0005, **** p< 0.0001.

